# Identifying parsimonious pathways of accumulation and convergent evolution from binary data

**DOI:** 10.1101/2024.11.06.622201

**Authors:** Konstantinos Giannakis, Olav N. L. Aga, Marcus T. Moen, Pål G. Drange, Iain G. Johnston

## Abstract

How stereotypical, and hence predictable, are evolutionary and accumulation dynamics? Here we consider processes – from genome evolution to cancer progression – involving the irreversible accumulation of binary features (characters), which can be modelled as Markov processes on a hypercubic transition network. We seek subgraphs of such networks that can generate a given set of paired before-after observations and minimize a topological cost function, involving criteria on out-branching which are interpretable in terms of biological parsimony. A transition network supporting a single, deterministic dynamic pathway is maximally simple and lowest cost, and branches (corresponding to possibly different next steps) increase cost, particularly if these branches are “deep”, occurring at early stages in the dynamics. In this sense, the lowest-cost subgraph measures how stereotypical the evolutionary or accumulation process is, and also identifies good start points for likelihood-based inference. The problem is solvable in polynomial time for cross-sectional observations by building on an existing method due to Gutin, and we provide a polynomial-time estimate in the more general case of pairs of observed states. We use this approach to define a “stereotypy index” reflecting the extent of evolutionary predictability. We demonstrate use cases in the evolution of antimicrobial resistance, organelle genomes, and cancer progression, and provide a software implementation at https://github.com/StochasticBiology/hyperDAGs.

## Introduction

Many processes in evolutionary biology, disease progression, and beyond can be viewed as the coupled acquisition of binary features over time. In evolutionary biology, these features may be binary characters acquired by species on a phylogeny. In cancer progression, the features may be presence/absence markers of particular mutations in a developing tumour. Other instances of such “accumulation models” include the buildup of symptoms in patients through disease progression, the progress of students through online courses, pathogens evolving resistance to different drugs, and more (Beerenwinkel et al., 2005; Diaz-Uriarte, 2023; Diaz-Uriarte & Herrera-Nieto, 2022; Johnston et al., 2019; Johnston & Røyrvik, 2020; Peach et al., 2021; Renz et al., 2024; Schill et al., 2024).

Several approaches have been developed for inferring the dynamic details of these processes from observed data. Foundational phylogenetic work by Pagel (Pagel, 1994) considered inferring the coevolutionary connection of two binary characters on a phylogeny. More recent work in the phylogenetic literature has expanded this picture to larger sets of binary characters (Boyko & Beaulieu, 2021) and to combined discrete and continuous characters (Boyko et al., 2023). In parallel, work motivated by cancer progression (where samples are often not phylogenetically connected) has developed a wide set of approaches, typically more focused on the features and interactions between them than on the particular states (Beerenwinkel et al., 2015; Diaz-Uriarte, 2023; Diaz-Uriarte & Herrera-Nieto, 2022; Schill et al., 2024; Schwartz & SchäEer, 2017). These approaches may be based on logical dependencies between acquisition features (Desper et al., 1999; Szabo & Boucher, 2008; Zhang & Wang, 2018), including approaches using conjunctive and disjunctive Bayesian networks (Angaroni et al., 2022; Beerenwinkel et al., 2007; Gerstung et al., 2009; Montazeri et al., 2016; Nicol et al., 2021), permutation analysis (Peterson & Kovyrshina, 2017), or stochastic influences between feature acquisitions (Hjelm et al., 2006; Schill et al., 2020).

Bridges between phylogenetically-embedded data and accumulation models have been developed. A class of models under the umbrella of “hypercubic inference” allow both cross-sectional and phylogenetically embedded data: these include HyperTraPS (Aga et al., 2024; Greenbury et al., 2020; Johnston & Williams, 2016), HyperHMM (Dauda et al., 2024; Moen & Johnston, 2023), and a recent approach employing the Mk model (Johnston & Diaz-Uriarte, 2024). The cancer field, particularly catalysed by the emergence of single-cell data, has developed methods for inferring dynamics of “phylogenetically” embedded cancer progression data (trees corresponding to patterns of cellular descent) (Luo et al., 2023; Ross & Markowetz, 2016).

There is a less-explored parallel to the inference problems that have been studied here to date. These approaches explore, explicitly or implicitly, different weights on a transition network linking states, typically attempting to maximise the likelihood associated with a dataset by varying transition weights. In these contexts, the dataset can be represented as a set of before-after pairs of states (Fig. 1). In the case of independent, cross-sectional observations, the “before” state is always a precursor with no features, and each “after” state is an independent observation (Fig. 1C). For longitudinal or phylogenetically linked observations, the “before” states are ancestral states and the “after” states are descendant states throughout the lineages involved, with unobserved states typically being reconstructed or inferred (Fig. 1D). Given such data, we ask instead: what is the topologically most parsimonious model of evolutionary dynamics that has a nonzero likelihood? In other words, what is the simplest set of nonzero edges in the transition network so that all observations are covered?

**Figure 1.**
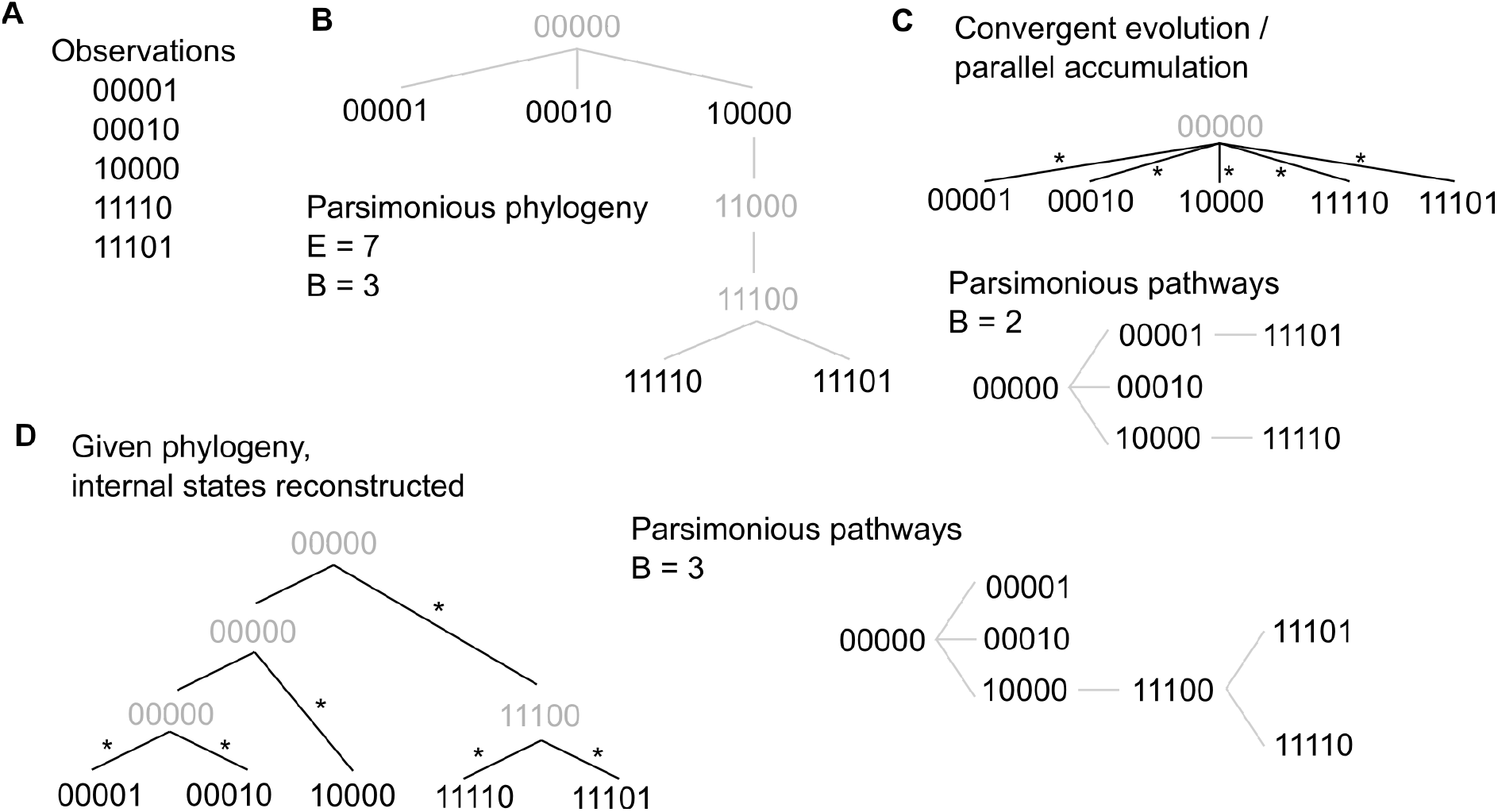
Parsimonious phylogenies vs parsimonious evolutionary pathways. Given a set of observations of L = 5 binary characters (A), a common task is to construct a parsimonious phylogeny that links the observations while requiring the fewest edges or independent character changes (B). In (B), a minimal phylogeny has E = 7 edges and B = 3 branches in excess of a single lineage (Eqn. 2). By contrast, in accumulation modelling, the relationship between observations is typically assumed (C-D). In convergent evolution / parallel accumulation (C), each observation reflects an independent progression from a precursor state (assumed to be 00000), and so reflects an independent before-after pair of “observations” of the evolutionary process (labelled with stars). Our goal is to find parsimonious sets of evolutionary pathways that capture each before-after observation while minimising the number of separate branches, or different routes evolution can take. We may also have an independently estimated phylogeny with reconstructed internal states (D). This will change the set of before-after “observations” (again, each edge with a star) and hence the parsimonious pathway set.

This is a different view of “parsimony” to that often considered in evolutionary (and cancer progression) work. From the perspective of reconstructing phylogenies, evolution is typically viewed as a collection of rare, independent events. The most parsimonious explanation for a set of observations is therefore the one that minimises the number of events required to explain those observations (Fig. 1A-B). In convergent evolution, the same event can occur many times independently (Fig. 1C) – as in famous cases like the camera eye (in cephalopods, vertebrates, and cnidaria), and C_4_ photosynthesis (evolved independently in over 60 lineages (Williams et al., 2013)). Of course, many cases outside evolutionary biology – cancer progression in independent patients, for example – also fall within this category of multiple parallel lineages independently accumulating features.

For our question, the target of our estimation is not a parsimonious phylogeny connecting observations (Fig. 1B), for which many approaches exist for discrete characters (Felsenstein, 2003), including Camin-Sokal, Wagner, and Dollo criteria for parsimony (Camin & Sokal, 1965; Day et al., 1986; Farris, 1970, 1977). Indeed, the “phylogeny” is just a collection of independent stumps in the case of cross-sectional data, or a star graph in the case of convergent evolution (Fig. 1C); in other accumulation modelling contexts we normally take a phylogeny as a given input (Fig. 1D). Instead we are interested in identifying a parsimonious set of “pathways” that evolution can take – that is, the alternative orderings in which the set of traits can be acquired. In this picture, each independent lineage proceeds down a particular pathway (Fig. 1C-D). The most parsimonious pathway set compatible with observations is then of both scientific and technical interest. Scientifically, the simplest evolutionary model corresponds to the most deterministic picture of evolution – and hence measures how stereotypical (and hence predictable) the progression of a given process is. For example, if all observations can be captured by a single model pathway with no branching, the accumulation process is completely deterministic. A single branch means that evolution always proceeds down one of two pathways, and so on. Technically, the simplest model may serve as a good initial guess for the parameter searches in the likelihood-based approaches above, which often spend considerable time converging to a relevant region of parameter space.

We proceed by introducing some definitions of use for this work. We then introduce how an algorithm from (Bang-Jensen & Gutin, 2018; Gutin et al., 2008) can be adapted to solve this problem for cross-sectional observations, and introduce a new approach building upon this picture for the more general cases of before-after pairs of observations.

## Methods

We begin by defining some properties of graphs that will be useful in describing accumulation of binary traits. Fig. 2 illustrates some of the core concepts for this study.

**Figure 2.**
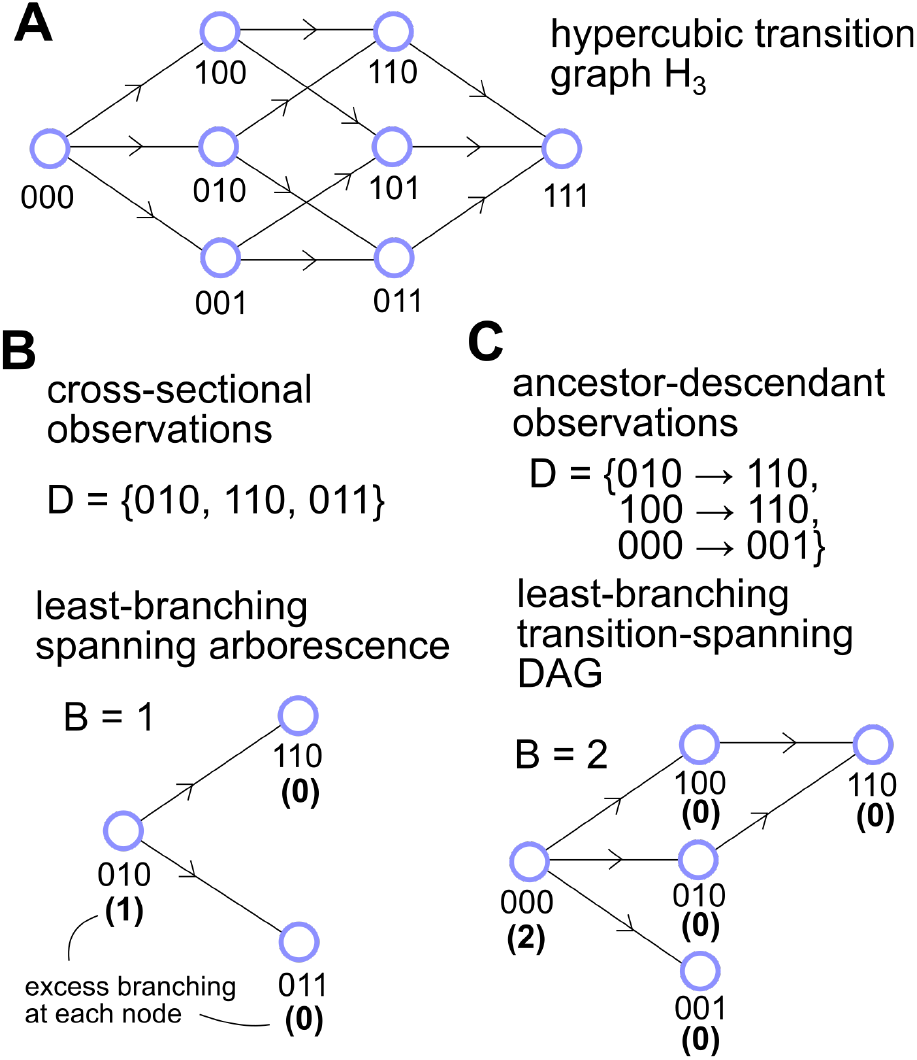
Concepts and definitions. (A) A hypercubic transition graph – a DAG capturing possible transitions between states described by binary strings. Here the H_3_ graph captures a system with L=3 binary features; the layer L(*v*) increases from 0 to 3 from left to right. (B) Given a set of cross-sectional observations *D*, a least-branching spanning arborescence is a subgraph of the hypercubic transition graph containing all states in *D* and minimising excess branching *B* (Eqn. 2). *B* is the sum of excess branching values over all nodes (in brackets). (C) Given a dataset consisting of paired ancestor-descendant observations, a least-branching transition-spanning DAG (LBTSD) is a subgraph with minimal excess branching that has a path linking every ancestor-descendant pair. Here, (B) and (C) are also the simplest spanning arborescence and simplest transition-spanning DAG respectively (Definition 9), as we cannot move the branching edges to increase the layer sum of the subgraphs (Eqn. 3).

### Definition 1

Hypercubic distance between binary strings *s*_*1*_ and *s*_*2*_. Define *d*(*s*_*1*_, *s*_*2*_) = Hamming(*s*_*1*_, *s*_*2*_) if *s*_*1i*_ <= *s*_*2i*_ for all *i*; d(*s*_*1*_, *s*_*2*_) = ∞ otherwise.

For example, d(100, 110) = 1 (one acquisition gets us from 100 to 110), d(100, 000) = ∞ (no set of acquisitions can get us from 100 to 000).

### Definition 2

Hypercubic transition graph and layers. Let *B*^*L*^ be the set of binary strings of length *L*. Let *H*_*L*_ be the unique hypercubic transition graph for *L* characters, that is the DAG (directed acyclic graph) where *V*(*H*_*L*_) = *B*^*L*^ and *E*(*H*_*L*_) = {*i* → *j*: *d*(*i, j*) =1} (Fig. 2A). The root of *H*_*L*_ is the node of all zeroes, which we write *0*^*L*^ for brevity. All paths on *H*_*L*_ eventually reach the node of all ones, which we write *1*^*L*^ for brevity.

Given a hypercubic transition graph, the layer L(*v*) of a vertex *v* is the number of 1s in the binary string label for *v* (Fig. 2A), and the layer L(*e*) of an edge *e* is the layer of its source vertex.

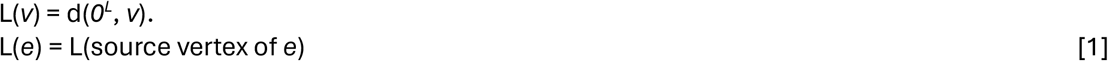

### Definition 3

Excess branching. Define excess branching *B*(*G*) for digraph *G* as

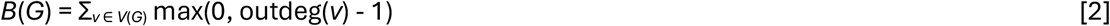

The max function is to account for the fact that leaves in the graphs have out-degree 0.

### Lemma 1

In an arborescence (directed tree), every branch creates a new leaf, so *B*(*G*) = |leaves(*G*)| - 1. If our DAG is not an arborescence, multiple branches may “resolve” to the same vertex and this relationship will not hold.

### Definition 4

Spanning arborescence. Given a dataset of nodes {*D*_*i*_} ∈ V(*H*_*L*_), a spanning arborescence for *D* is an arborescence *G* ∈ *H*_*L*_ such that {*D*_*i*_} ∈ V(G).

### Definition 5

Least branching spanning arborescence (LBSA). If *G* is a spanning arborescence for *D* such that *B*(*G*) (Eqn. 2) takes a minimum value over all spanning arborescences for *D, G* is a least branching spanning arborescence for *D* (Fig. 2B).

By Lemma 1, the problem of finding a least branching spanning arborescence for *D* is the same as finding a spanning arborescence with minimum leaf count. This is called (amongst other things) the minimum leaf out-branching problem (“out-branching” is used synonymously with “arborescence” in different fields). In general it is NP-hard, but for the arborescence structures we consider there is a polynomial solution, specifically because the problem is embedded in a directed acyclic graph (DAG). This solution, described in (Gutin et al., 2008) and detailed in (Bang-Jensen & Gutin, 2018), will be referred to as “Gutin’s algorithm”.

### Definition 6

Transition-spanning DAG. Let D be a dataset consisting of pairs {*D*_*1i*_, *D*_*2i*_}. If *G* is a directed acyclic graph such that:

1. *G* ∈ *H*_*L*_, *D*_*1i*_ ∈ V(*G*) and *D*_*2i*_ ∈ V(*G*) for all *i;*
2. A path exists in *G* from *D*_*1i*_ to *D*_*2i*_ for all *i*,
3. A path exists in G from *0*^*L*^ to every *D*_*1i*_ and every *D*_*2i*_ for all *i*, then *G* is a transition-spanning DAG for *D*.

A transition-spanning DAG has stricter requirements than a spanning arborescence because we require a specific set of paths (between pairs of vertices), and a path to the “precursor” node *0*^*L*^ for every vertex to be included. The reason for (c) is that in accumulation modelling we typically assume that either the common ancestor (for phylogenetic observations) or the initial state (for cross-sectional or longitudinal observations) has none of the features under study (for example, no cancer-related mutations), so to describe the dynamics completely we require a path from *0*^*L*^ to every “ancestral” state (Fig. 1D).

### Definition 7

Least branching transition-spanning DAG (LBTSD). If *G* is a transition-spanning DAG for *D* such that *B*(*G*) (Eqn. 2) takes a minimum value over all transition-spanning DAGs for *D, G* is a least branching transition-spanning DAG for *D* (Fig. 2C).

Identifying the LBTSD informs about the smallest number of distinct evolutionary pathways that can account for all snapshot or longitudinal observations in a given dataset. The magnitude of *B*(*G*)+1 is the number of distinct evolutionary paths required: *B*(*G*) =0 means that a single, linear, deterministic progression of states can explain all observations; *B*(*G*) =1 means that the evolution of a given lineage proceeds down one of two pathways, and so on.

### Lemma 2

The identification of an LBTSD is NP-complete. See Appendix for proof, which provides a mapping from “Vertex Cover”, a known NP-complete problem (Garey & Johnson, 1979; Karp, 2010), to LBSTD identification.

### Definition 8

Layer sum. Given a subset of a hypercubic transition graph, the “layer sum” LS is the sum of layers (Eqn. 1) of edges that contribute to excess branching:

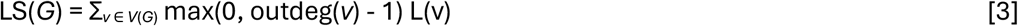

### Definition 9

Simplest spanning arborescence (SSA) and simplest transition-spanning DAG (STSD). A simplest spanning arborescence or transition-spanning DAG for data *D* is an LBSA or LBTSD for *D* where the layer sum LS (Eqn. 3) is maximised.

The SSA and STSD fulfil the criteria of having minimal evolutionary branching, and also ensure that this branching occurs as late as possible in the evolutionary process, so that the early dynamics of the system are as canalized and stereotypical as possible.

A “stereotypicality index” S can be defined for an STSD and its source dataset, reporting to what extent the inferred evolutionary or progression process is reproducible and predictable. A simple value for this index is

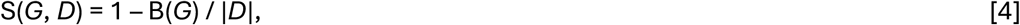

where B(*G*) is the branching count of the graph *G* and |*D*| is the number of observations in the dataset. Hence, if B(*G*) = 0 and all observations can be explained by a single evolutionary pathway, S = 1 for maximum stereotypy. If every observation in *D* requires a new branch to explain it, S → 0 for minimum stereotypy.

In many cases, the same transition occurs multiple times independently in a set of observations. Although such repeats do not affect the algorithms below, we propose using the original size of the dataset for |*D*|, as (intuitively) repeated independent observations imply more stereotypical dynamics in the underlying process. The statistic S can then be viewed as a readout of how “convergent” the evolutionary process is (qualitatively similar to, for example, the Wheatsheaf Index (Arbuckle et al., 2014)).

## Results

### Identifying a simplest spanning arborescence for cross-sectional data

In the cross-sectional case, we have a set of independent observations and require a subgraph of the hypercubic transition graph that hits each corresponding vertex. If we picture the ancestral case (initial conditions of the system) as *0*^*L*^, this is equivalent to the ancestor-descendant picture where every ancestor is *0*^*L*^. In that picture, any optimal transition-spanning DAG is intuitively an optimal spanning arborescence rooted at *0*^*L*^, because any node with multiple incoming paths from the root represents a redundancy that can be removed (with corresponding decrease in *B*(*G*)). A solution for this can be obtained in polynomial time with Gutin’s algorithm (Bang-Jensen & Gutin, 2018; Gutin et al., 2008).

Given such an LBSA, an estimate for the simplest spanning arborescence (SSA) can be constructed by “rewiring” the graph in such a way as to preserve excess branching (Eqn. 2) while shifting the source nodes of branching as far as possible from the root node *0*^*L*^ (Fig. 3A-B).

**Figure 3.**
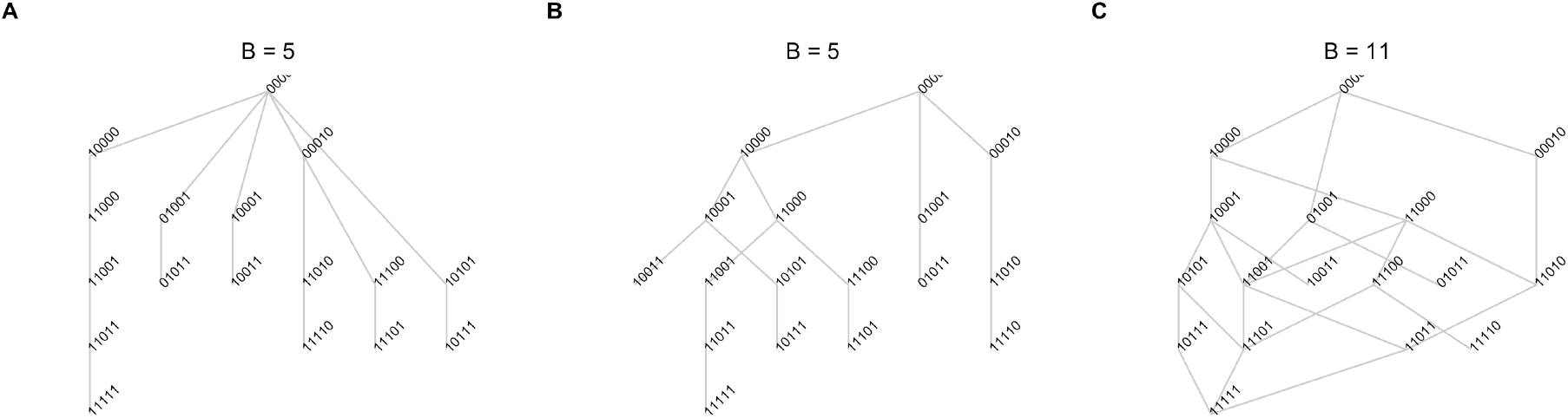
Outputs at different stages of Algorithm 1 and 2. (A) LBSA (least branching spanning arborescence) computed with Gutin’s algorithm for an example cross-sectional *L* = 5 dataset with 427 transitions (54 unique) involving 17 unique states. (B) Estimated SSA (simplest spanning arborescence): the LBSA rewired with Algorithm 1 to compute an SSA, with the additional minimisation condition on layer sum bringing branch points “lower” in the arborescence (so the evolutionary process remains as constrained as possible for as long as possible). Both (A) and (B) have excess branching B(*G*) = 5; layer sum LS(*G*) = 0 for (A) and 5 for (B). (C) Estimated STSD (simplest transition-spanning DAG): Algorithm 2 applied to paired ancestor-descendant transitions in the same dataset, with *B* = 11. Stereotypy S = 0.97.

#### Algorithm 1 Rewiring an LBSA to estimate an SSA

Takes input data {*D*}.

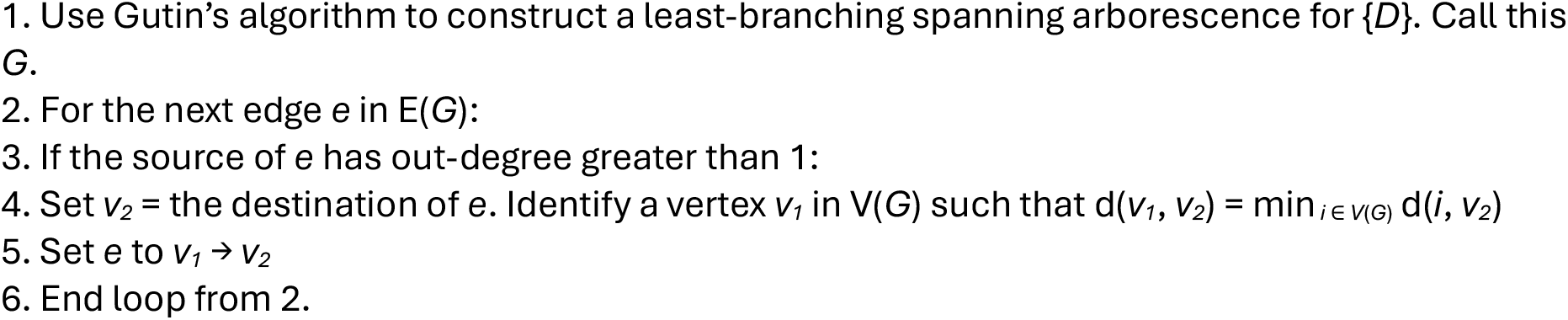

Algorithm 1 then takes an LBSA and shifts the sources of branches in the arborescence to vertices that are as close as possible to the destination vertex (and hence as far from *0*^*L*^ as possible). This preserves B(*G*) while increasing the layer sum LS(*G*), respecting the criteria for the SSA.

### Identifying a simplest transition-spanning DAG for ancestor-descendant data

In the longitudinal case, the data may require several different ancestors for a given descendant, and the solution will not in general be an arborescence but is a DAG (Definitions 7, 9). By Lemma 2 (see Appendix), the identification of the least-branching transition-spanning DAG is NP-complete, and so we work towards estimating good solutions rather than identifying a global optimum.

#### Algorithm 2 Estimating the STSD

Takes input data {*D*_*1i*_, *D*_*2i*_}.

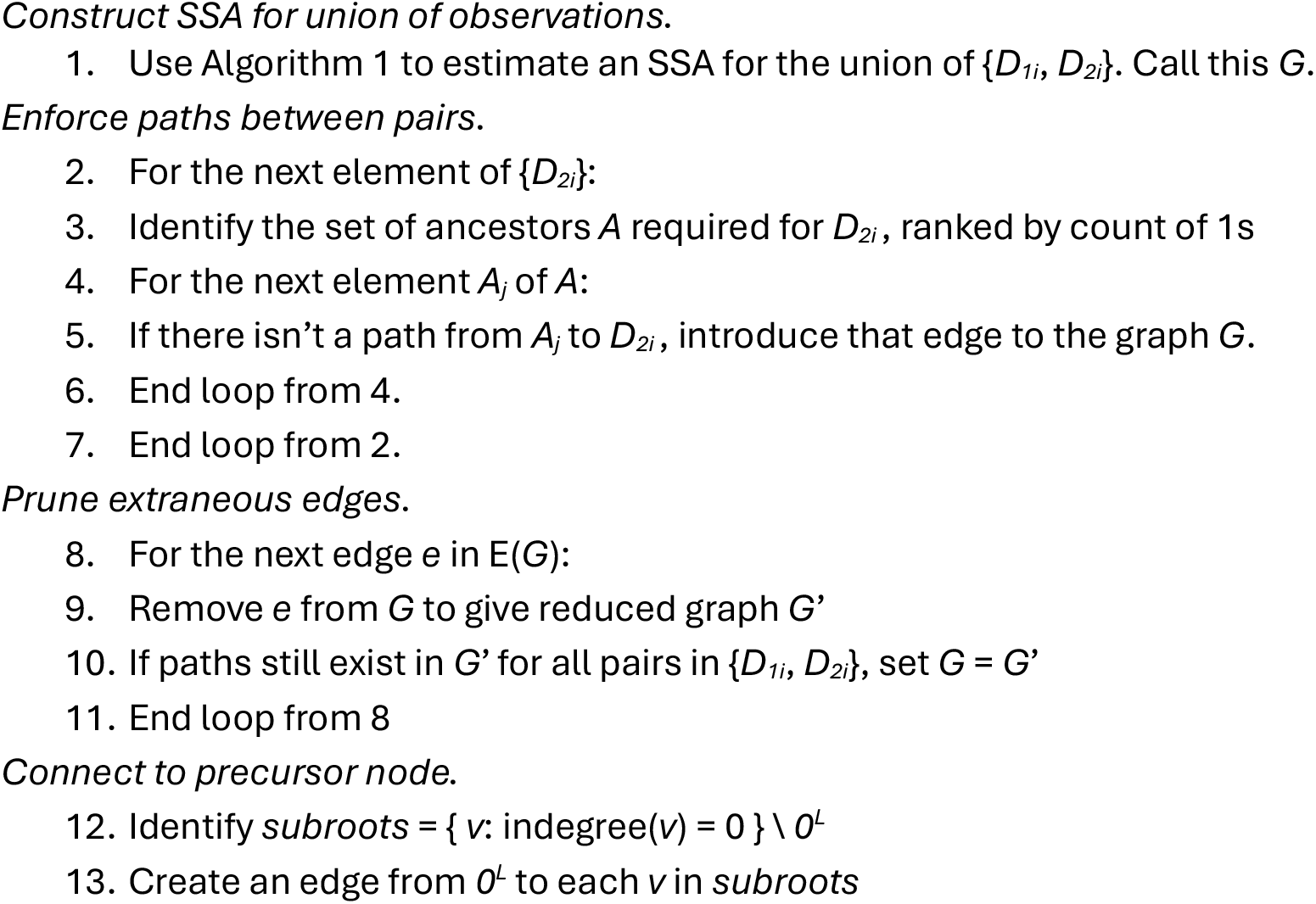

Algorithm 2 first builds an minimal arborescence hitting all states in the dataset (forgetting ancestor-descendant relationships). It then introduces additional edges required to fulfil these relationships, working case-by-case. In general the arborescence can become a DAG in this step. It then prunes edges from the expanded graph if they are not necessary to retain all transitions in the dataset (for example, if the case-by-case expansion introduced redundant edges). Finally, it ensures there is a path from *0*^*L*^ to every vertex. An example of progress through the different stages of this approach, for an L = 5 dataset, is shown in Fig. 3.

### Example: anti-microbial resistance in tuberculosis

We will first demonstrate the outputs of Algorithms 1 and 2 on a dataset on the evolution of drug resistance in tuberculosis (Casali et al., 2014). Here, several hundred isolates of tuberculosis were sampled from Russia, and each isolate’s susceptibility or resistance (0 or 1 respectively) to each of 10 drugs was recorded. The phylogeny linking these isolates was constructed in the original study, and previous work (Greenbury et al., 2020) has computed the set of ancestor-descendant transitions corresponding to these observations.

In Fig. 3, the dataset used was a subset of these data involving only the first 5 drugs, so that our binary features correspond to resistance to INH (isoniazid); RIF (rifampicin, rifampin in the United States); PZA (pyrazinamide); EMB (ethambutol); and STR (streptomycin).

The full dataset, involving resistance profiles to each of *L* = 10 drugs (INH; RIF; PZA; EMB; STR as before; additionally AMI (amikacin); CAP (capreomycin); MOX (moxifloxacin); OFL (ofloxacin); and PRO (prothionamide)), can also readily be analysed with this approach (Fig. 4). In both cases, several aspects of the subgraphs identified agree with the findings of previous work characterising evolutionary dynamics in this particular case study (Aga et al., 2024; Greenbury et al., 2020; Moen & Johnston, 2023). In these approaches, the first two features are typically inferred to be acquired first, agreeing with the large set of paths descending from the 11000… state (equivalent to the feature set {INF, RIF}) in Fig. 4A (and 3C). The {INF, RIF, STR} state and its immediate descendant {INF, RIF, EMB, STR} have the highest out-degrees in the whole network, reflecting that many observations can be considered descendants of these states. A parsimonious evolutionary model would include these states as likely early stages, and indeed these states are those that appear with highest probability in both maximum likelihood (Moen & Johnston, 2023) and fully Bayesian approaches (Aga et al., 2024; Greenbury et al., 2020) inferring probabilistic dynamics in this system. The subsequent more extensive branching also aligns with the multiple different pathways supported in the full inference picture.

**Figure 4.**
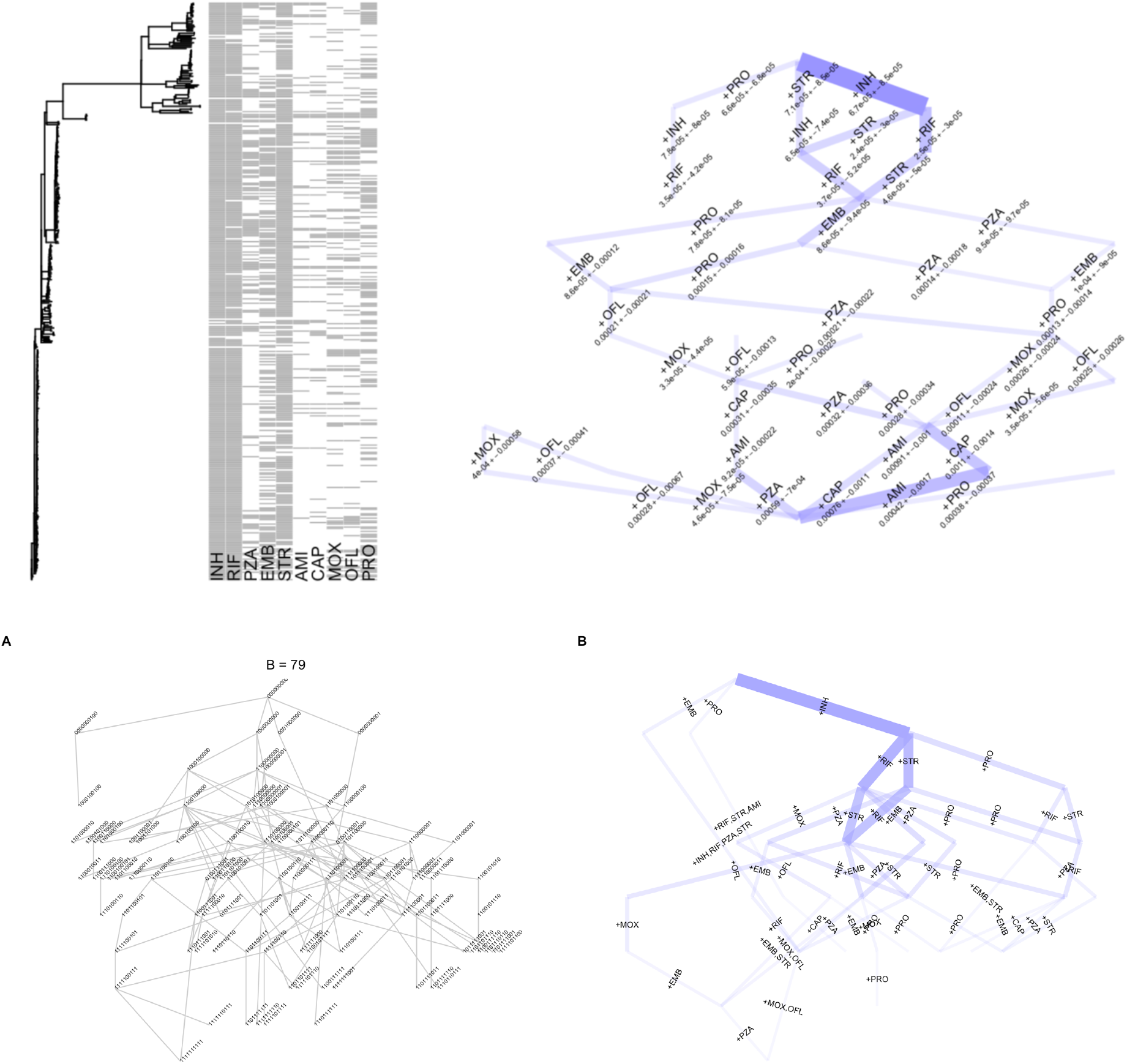
Evolution of multidrug resistance in tuberculosis, and connection to inference of dynamics. (top) Tuberculosis dataset and accumulation dynamics inferred using HyperTraPS-CT, from (Aga et al., 2024). (A) Algorithm 2 applied to the full tuberculosis dataset with *L* = 10 features and 427 transitions (210 unique) involving 99 unique states. (B) The same network, with edge thickness determined by out-neighbourhood size (a proxy for evolutionary importance), and edges under a threshold of 5 pruned. Edge labels give the specific drug resistance(s) acquired in each transition. Stereotypy S = 0.81.

We can also demonstrate how the purely topological characteristics of these inferred graphs can provide a fast first approximation to some aspects of the full dynamic inference. To quantify the reasoning above, where states with many descendants are likely to be parsimonious evolutionary precursors, we calculate the out-neighbourhood of each node, defined as the number of nodes which can be reached from that node. Labelling the edges in the inferred graph by the size of this out-neighbourhood then reflects its importance in a parsimonious evolutionary explanation of the data, which is related to how likely it is to appear in evolutionary dynamics under such a model (Fig. 4B). This picture, of course, does not take into account the relative observation frequencies of different states or transitions and has no underlying probabilistic model, just an estimation based on parsimonious topological characteristics.

### Example: cancer progression

Cancer progression often involves the accumulation of different mutations over time. For a comparison with several recent approaches, we use the single-cell genomic dataset of acute myeloid leukemia evolution from (Morita et al., 2020). This dataset consists of a set of trees linking “ancestral” and “descendant” genomic observations in cancer patients, represented as barcodes describing the presence/absence of a given genetic mutation in a sample.

The structure of the graph in Fig. 5 reflects known properties from other inference approaches applied to these data (Aga et al., 2024; Luo et al., 2023). A rather broad set of initial steps correspond to the large range of single mutations observed throughout the dataset. No paths longer than length 8 exist, reflecting the maximum cardinality of a set of mutations found in any sample. Weighting edges by out-neighbourhood size as described above shows that the most common first gene to be mutated is inferred to be DNMT3, also observed in other inference-based approaches (Aga et al., 2024; Luo et al., 2023). The weights assigned to other initial steps are different from these approaches. HyperTraPS, for example, orders likely first steps DNMT3, FLT3, IDH2, NRAS, NPM1, TET2 (with TreeMHN largely agreeing). This topology-based approach places more weight on early TET2 acquisition, likely because the set of possible descendants for {TET2} and the set of possible descendants for {DNMT3} are more disjoint than for other descendant sets. Following {TET2}, the most likely next acquisition is inferred to be DNMT3 across approaches; the topology approach has TET2 a reciprocal likely immediate followup to {DNMT3}, while HyperTraPS does not place TET2 as a likely next step after {DNMT3} (though IDH2, NPM1, and other followup acquisitions are highlighted in both approaches). The common acquisition of DNMT3 after alternative first steps is also observed across all approaches.

**Figure 5.**
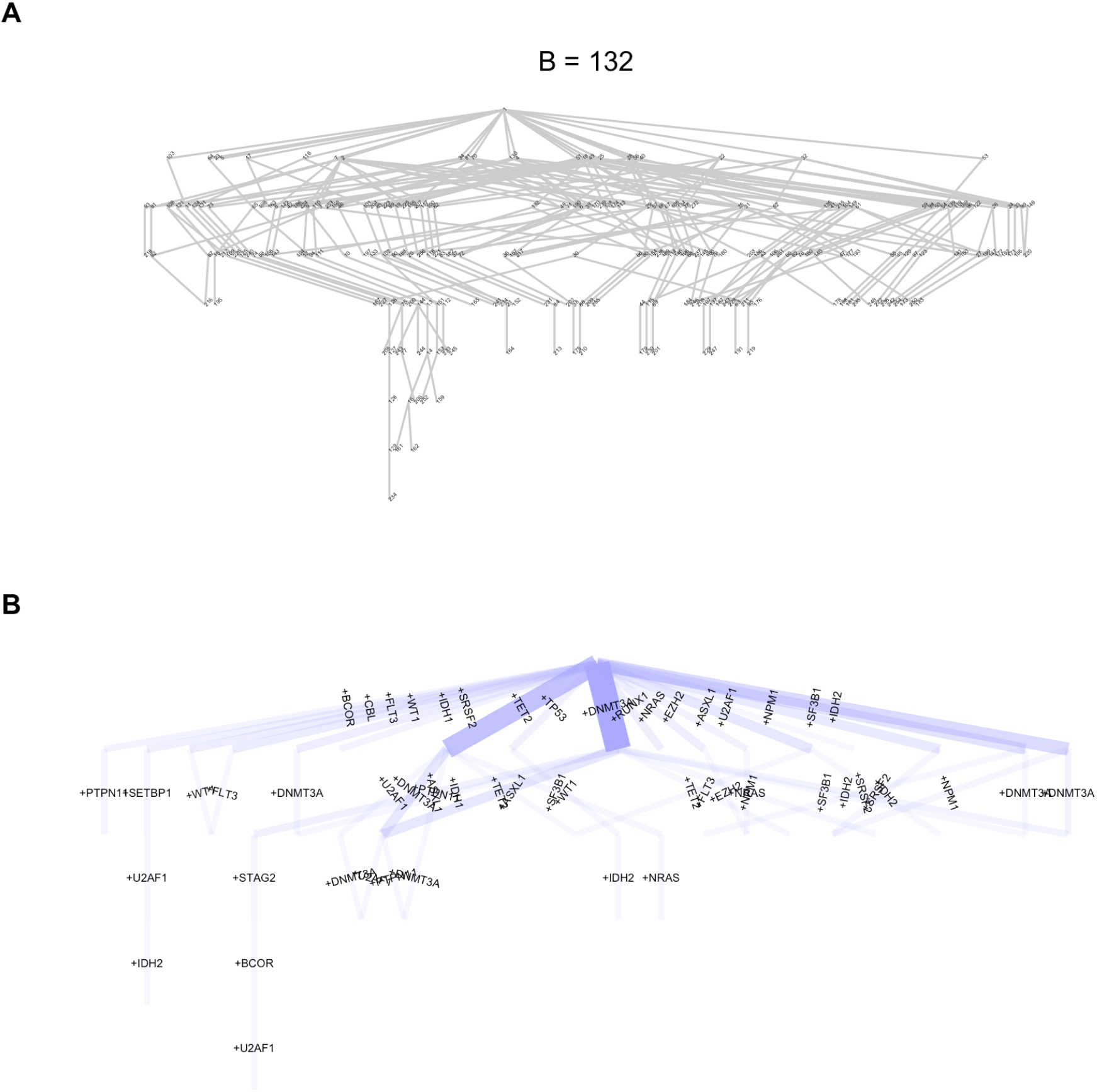
Pathways of single-cell cancer evolution. (A) Algorithm 2 applied to the acute myeloid leukemia dataset with *L* = 31 features and 465 transitions (278 unique) involving 256 unique states. (B) Graph weighted by out-neighbourhood size and pruned as in Fig. 4B. Stereotypy S = 0.72.

### Example: organelle gene loss

Mitochondria and chloroplasts are compartments in eukaryotic cells (the domain of life containing organisms that are not bacteria or archaea). They were originally independent organisms that were absorbed by ancestral eukaryotes billions of years ago, and since these “endosymbiotic” absorption events, their genomes have been dramatically reduced. From presumably thousands of original genes, only small numbers of protein-coding genes are retained in modern mitochondria (<70) and chloroplasts (<250). The pathways by which these genes are lost – and why some genes are lost but not others -- is the subject of considerable debate and ongoing data-driven research (Allen, 2015; Björkholm et al., 2015; Butenko et al., 2024; Giannakis et al., 2022; Johnston & Williams, 2016; Keeling, 2010; Kelly, 2021; Lynch et al., 2006; Roger et al., 2017; Smith & Keeling, 2015; von Heijne, 1986).

Recent work exploring the patterns of organelle gene loss has either used intensive computational resource for the mitochondrial case (Johnston & Williams, 2016) or considered only summary statistics, not full evolutionary spaces, for both organelles (Giannakis et al., 2022). The approach in this article allows the first characterisation of plausible hypercube-embedded loss pathways based on large-scale genomic data (Fig. 6). We used data on gene presence/absence from NCBI’s organelle genome database (O’Leary et al., 2016), curated via a pipeline in (Giannakis et al., 2022), which attempts to resolve inconsistencies in gene labelling by clustering sequences by genetic similarity and iteratively relabelling the identity of these clusters until consensus is reached (the outputs are also compared to manual relabelling of the diverse gene names in the dataset). The converged set of gene presence/absence markers are then combined with an estimated phylogeny to identify transitions involving loss of organelle genes throughout eukaryotic history. The processed dataset then contains before-after observations with presence or absence markers for each gene in the superset of organelle-encoded protein-coding genes found across eukaryotes.

**Figure 6.**
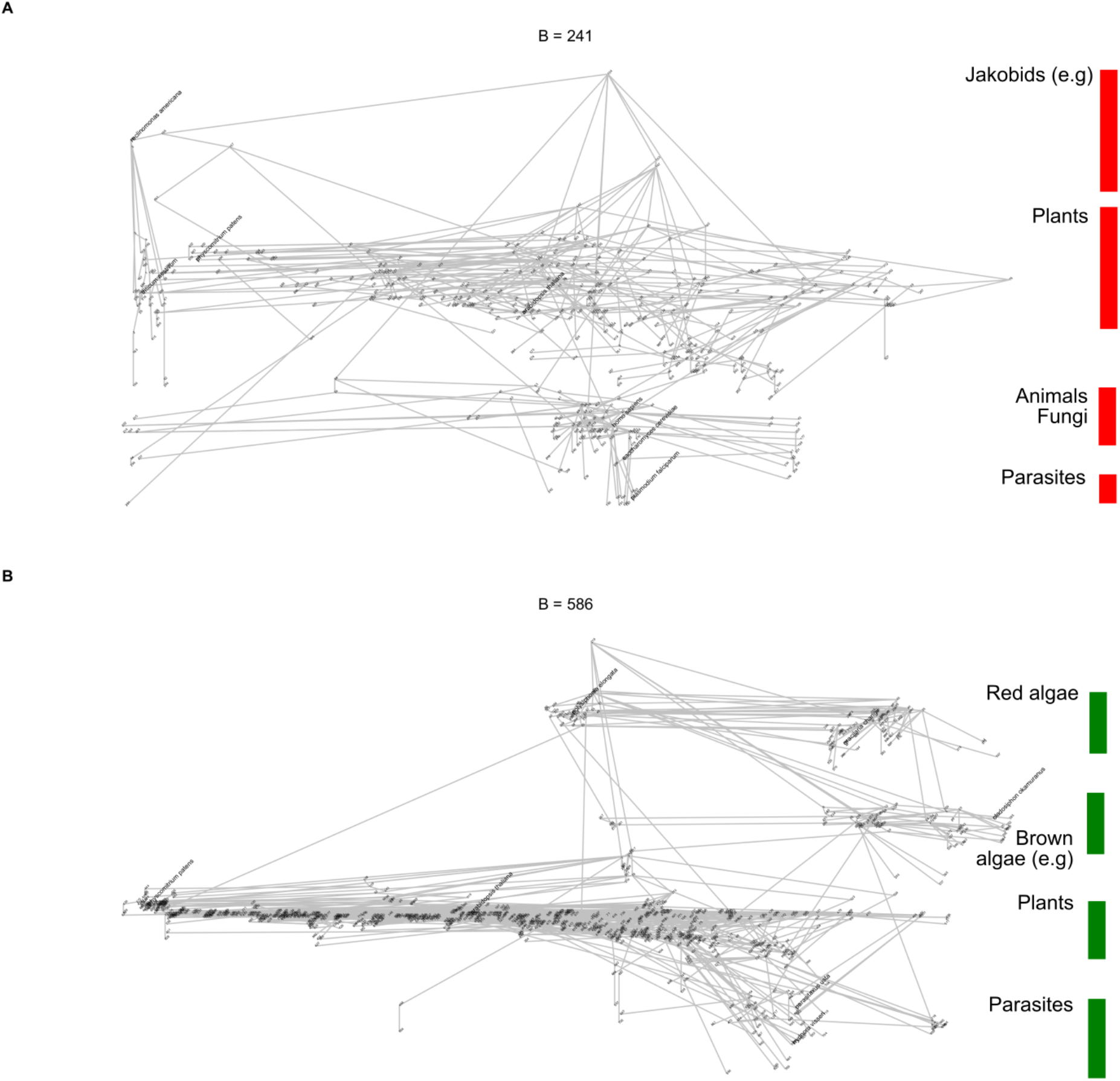
Mitochondrial and plastid gene loss. (A) Mitochondrial DNA (mtDNA), *L* = 66, with 924 transitions (494 unique) involving 420 unique states. Stereotypy S = 0.74. (B) Plastid DNA (ptDNA), *L* = 197, with 3065 transitions (1119 unique) involving 937 unique states (ptDNA). Clusters corresponding to different eukaryotic organisms are labelled with vertical bars. Stereotypy S = 0.81. Some example species profiles are labelled at their corresponding nodes in both cases; vertical sidebars give some broad types of organism that are represented in the various layer clusters of the graphs.

The number of features in these datasets is large enough that inference using the approaches mentioned above is computationally challenging (though not intractable with some methods including HyperTraPS (Aga et al., 2024; Johnston & Williams, 2016)). This topological approach, on the other hand, proceeds quickly and these cases are very tractable on a single core.

The results of Algorithm 2 are shown in Fig. 6. Biological insight is available from the structure of these graphs. In the mtDNA case, different domains of life are clearly visible as different layers of the hypercubic graphs. Jakobids like *Reclinomonas americana*, retaining the highest counts of mtDNA genes (Burger et al., 2013), are early branches from the root of the tree. A wide set of branching trajectories in the central layers of the structure correspond to plants, which retain moderate complements of mtDNA genes and are known for the plasticity, diversity, and dynamism of the mtDNA genes they retain (Johnston, 2019; Sloan et al., 2012, 2017). A narrower cluster of limited branching at lower layers corresponds to animals and fungi, which exhibit smaller and much less diverse complements of mtDNA genes. From this limited set, the lowest-layer transitions run to states corresponding to parasitic organisms like Apicomplexans, which retain only 2-3 protein-coding genes (Hjort et al., 2010; Makiuchi & Nozaki, 2014; Oborník & Lukeš, 2015).

The structure of the ptDNA graph is rather different. A dramatic early branch separates two clusters at high layers. This corresponds to deep-branching evolutionary lineages with different histories of plastid evolution – Rhodophyta (red algae) and other organisms (Keeling, 2010). The majority of nodes fall within a very restricted set of layers further down the hypercube: many different gene profiles, but with very similar overall gene counts. This large set corresponds to plants, where there is rather less diversity in gene profiles in ptDNA compared to mtDNA. Some descending transitions lead to rather more reduced ptDNA profiles – again, these correspond to parasitic organisms, whose lifestyle has led to the atrophy of photosynthetic machinery (Fichera & Roos, 1997; Krause, 2008).

## Discussion

We have introduced an approach for estimating the evolutionarily simplest explanation for a given accumulation system. The concept of evolutionary simplicity here refers to minimizing the number of “choices” a system faces as it evolves through time – the number of different pathways the dynamics could follow. Quantifying these as branches in a transition network allows the scaleable estimation of simple generative mechanisms and a consequent quantification of the degree to which evolution is stereotypical and predictable.

We have not considered uncertainty in observations, instead assuming that all observed transitions are perfectly observed. Uncertain features in observations would mean that a set of vertices, rather than a single vertex, could be compatible with a given observed state, and the algorithms we use do not have the capacity to search over such sets for an optimal picture. This represents a potentially valuable future step for this research, as measurement uncertainty features in many accumulation modelling pictures. We have also assumed a complete absence of irreversibility, so that once acquired, a feature is never lost. This picture is appropriate for some accumulation models but will neglect potentially important dynamics in other cases. Including reversible transitions is possible in principle but dramatically increases the complexity of the problem; the approach here would no longer be tractable in polynomial time, and parallel approaches for inference are also much more computationally demanding with reversibility (Johnston & Diaz-Uriarte, 2024).

Our approach identifies a single estimate for a problem that is likely degenerate: in general several LBSAs, SSA, and LBTSDs will exist with the same branching and layer count statistics. Further, although Gutin’s algorithm (Bang-Jensen & Gutin, 2018; Gutin et al., 2008) guarantees that the LBSA identified will be optimal, the followup approaches in the algorithms we describe here do not guarantee optimality. The “rewiring” to minimize layer sum will not increase branching count, but more optimal methods for introducing and pruning edges corresponding to transitions may be possible, and approaches to identify not just a single estimate but the full set of degenerate estimates may follow. However, this method already has desirable properties of (to our knowledge) novelty, performance, biological insight, and links with inference approaches.

Several related questions have been studied throughout both the phylogenetic and the informatics literature. First, the aforementioned task of reconstructing parsimonious phylogenies (Camin & Sokal, 1965; Day et al., 1986; Farris, 1970, 1977; Felsenstein, 2003), including “perfect phylogenies” (every character change occurs only once, also called the Steiner Tree problem on phylogenies (Fernández-Baca, 2001)) and “near-perfect phylogenies” (Fernández-Baca & Lagergren, 2003; Sridhar et al., 2007) and minimum-evolution criteria (Catanzaro et al., 2015). These tasks are also considered in the cancer progression literature, where a phylogeny-like structure is inferred to describe a parsimonious history of tumour development compatible with observed mutational profiles: for example in (El-Kebir, 2018; Gao et al., 2022; Jahn et al., 2016). As described above, the task we address here is not reconstructing a phylogeny – which we take as given – assuming minimal evolutionary change, but rather reconstructing the dynamics of character evolution that occur on that given phylogeny when evolutionary change is viewed as common and repeatable. In graph theory, we have leant heavily on Gutin’s approach for identifying arborescences spanning points while minimisation out-branching (Bang-Jensen & Gutin, 2018; Gutin et al., 2008). Some similar questions involve finding minimal spanning trees with constraints on the number of leaves (Fernandes & Gouveia, 1998), arborescence optimisations based on edge properties (Georgiadis, 2003) (following classical work by Edmonds (Edmonds, 1967) and Chu & Liu (Chu, 1965)), and particularly minimal Steiner arborescences on directed hypercubes (Mahapatra et al., 2024), though these typically focus on properties of edges (like weights) as the criterion for minimisation, while we focus on out-branching.

Across the biological case studies we have considered, the stereotypy index is both high and constrained (between 0.7 and 0.85), suggesting that most transitions do not require individual evolutionary pathways to explain them. This further suggests a degree of “canalization” of the corresponding evolutionary and progressive dynamics, so that intrinsic or external features lead to the system evolving according to somewhat reproducible rules – or, in other words, a degree of convergence in the evolutionary processes (Giannakis et al., 2022, 2024; Williams et al., 2013). Two interesting questions arise for further investigation: first, how does this statistic for “degree of convergence” relate to alternatives (for example, the Wheatsheaf Index (Arbuckle et al., 2014; Speed & Arbuckle, 2017)) for instances where the two can naturally be compared? And second, what range can stereotypy index take across biological instances – in other words, which evolutionary processes are more or less predictable across life (Stayton, 2015)? We hope that further application of this approach to more biological systems will help explore these questions.

## Acknowledgments

This project has received funding from the European Research Council (ERC) under the European Union’s Horizon 2020 research and innovation program (grant agreement no. 805046 [EvoConBiO] to I.G.J.). This project was supported by the Trond Mohn Foundation (project HyperEvol under grant agreement no. TMS2021TMT09) through the Centre for Antimicrobial Resistance in Western Norway (CAMRIA) (TMS2020TMT11).

## Appendix

## Proof of Lemma 2

We aim to show that the identification of a least-branching transition-spanning DAG (Definition 7) is NP-complete; this is the reason why we therefore do not seek an exact algorithm. The problem can be defined formally as follows:

**Definition (excess branching)**. Given a directed graph *G*, and a vertex *ν*, we define *eb*_*G*_(*ν*) = max(deg_*G*_ (*ν*) − 1,0), i.e., the degree above 1. For the graph *G* itself, we define *eb*(*G*)—the excess branching of *G*—as

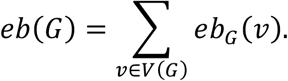

Given the definition of excess branching, the decision problem is then the following:

**Problem:** *Low Branching Connection Augmentation (LBCA)*

**Input:** a supergraph *H*^*∗*^, a set *R* of pairs of binary vectors, (*ν*_1_, *ν*_2_) corresponding to vertices in *H*^*∗*^ and a value *k* ∈ ℕ,

**Question:** Does there exist a subgraph *H* of *H*_*n*_ such that for each pair *ν*_1_, *ν*_2_, there is a path from *ν*_1_ to *ν*_2_ in *H* and such that the excess branching *eb*(*H*) ≤ *k*?

We will prove that the problem is NP-complete by reducing from *Vertex Cover* (Fig. A1). This problem is one of the original NP-complete problem by Karp, and we will provide a Karp reduction from *Vertex Cover* to our problem (Garey & Johnson, 1979; Karp, 2010).

Let (*G* = (*V, E*), *k*) be an instance of *Vertex Cover*. We will create an *equivalent* instance of our problem, where equivalent here means that the *Vertex Cover* instance is a yes-instance if and only if the input to our problem is a yes-instance. The reduction will run in polynomial time, and is hence a valid Karp reduction.

First, we reduce the problem to a problem where we do not demand finding a subgraph of *H*_*n*_, but a subgraph of a given DAG *H*^*∗*^, and then we will later see how to embed this into *H*_*n*_. For each vertex *ν* ∈ *V*, create a vertex *h*_*ν*_ ∈ *H*^*∗*^ and for each edge, we create a vertex *h*_*e*_ ∈ *H*^*∗*^. In addition, we create two terminal vertices *t*_0_ and *t*_1_ in *H*^*∗*^.

The purpose of *t*_0_ is that every vertex *h*_*ν*_ is connected to *t*_0_, and is required to remain connected to it in the subgraph. The purpose of *t*_1_ is that every vertex *h*_*e*_ will be required to be connected to *t*_1_ through some vertex *h*_*ν*_.

Then we add the following edges into *H*^*∗*^: If *e* = *uν* in *G*, add the two edges *eu* and *eν* into *H*^*∗*^. Finally, for every vertex *ν*, add the edge *νt*. This concludes the construction of the supergraph *H*^*∗*^.

The required pairs are simply

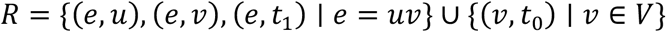

Now the input to the problem *Low Branching Connection Augmentation* is *H*^*∗*^, *R*, |*E*| + *k*.

**Lemma**. The input *G, k* is a yes-instance to *Vertex Cover* if and only if *H*^*∗*^, *R*, |*E*| + *k* is a yes-instance to *Low Branching Connection Augmentation*.

**Proof**. (⇒) Suppose that *G, k* is a yes-instance to *Vertex Cover* and let *C* be a solution of size at most *k*. Create a new graph *H* (Fig. A2) with vertex set *V*(*H*^*∗*^) and the following edge set: For each edge *e* = *uν* in *G*, keep the edges *h*_*e*_ *h*_*u*_ and *h*_*e*_ *h*_*ν*_.. For each vertex *ν* in *G*, keep the edge *h*_*ν*_. *t*_1_. Finally, for each vertex *ν* ∈ *C*, add the edge *h*_*ν*_*t*_1_. Clearly, *eb*(*H*) = |*E*| + |*C*| ≤ |*E*| + *k*, so the excess branching is within budget. Now, we trivially satisfy the requirements of the type (*h*_*e*_, *h*_*u*_) and (*h*_*ν*_., *t*_0_), since these are actual edges in *H*. Now, for the last type of requirement, let *e* = *uν* be an edge and (*h*_*e*_, *t*_1_) be a requirement. Since *C* is a vertex cover, at least one of *u* and *ν* are in *C*, and without loss of generality let *u* ∈ *C*. Then, by construction, *h*_*u*_ *t*_1_ is an edge, and hence (*h*_*u*_, *t*_1_ is satisfied through *h*_*u*_.

(⇐) Suppose that *H*^*∗*^, *R*, |*E*| + *k* is a yes-instance and let *H* be the solution of *eb*(*H*) ≤ |*E*| + *k*. Since every edge *e* in *G* must have an edge to two vertices, these edges account for |*E*| of the budget. Since every vertex *ν* in *G* is connected to *t*_0_, we have that all non-terminal vertices has degree at least 1. We conclude that there are *k* additional edges. Let *e* = *uν* be an edge in *G*. Then the vertex *h*_*e*_ has exactly two neighbors, namely *h*_*u*_ and *h*_*ν*_, and these are already accounted for, hence the only edges we can add are of type *h*_*ν*_*t*_1_ for vertices *ν* from *G*. Construct a set *C* = {*ν* ∈ *V*(*G*) ∣ *eb*(*h*_*ν*_) > 0}. We claim that |*C*| ≤ *k* and that *C* is a vertex cover in *G*. Since the remaining budget on the excess branching is *k*, we know that |*C*| ≤ *k*. Let *e* = *uν* be any edge from *G*. Since (*h*_*e*_, *t*_1_) ∈ *R*, and *H* is a valid solution, there is a path from *h*_*e*_ to *t*_1_ in *H*. The vertex *h*_*e*_ has only two neighbors, *h*_*u*_ and *h*_*ν*_, so at least one of *h*_*u*_ and *h*_*ν*_ has an edge to *t*_1_ in *H*, hence at least one of *u* or *ν* is in *C*. It follows that *C* is a vertex cover in *G*.

**Figure A1:**
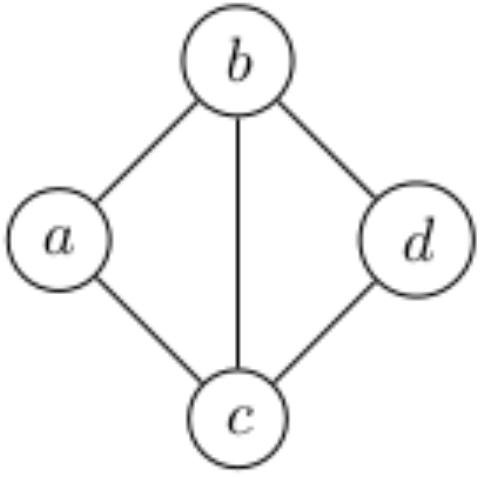
Input graph *G* = (*V, E*) with *k* = 2 to *Vertex Cover*. The unique *Vertex Cover* of size 2 is *C* = {*b, c*). It is easy to see that every edge is incident on *b* or *c* (or both).

**Figure A2:**
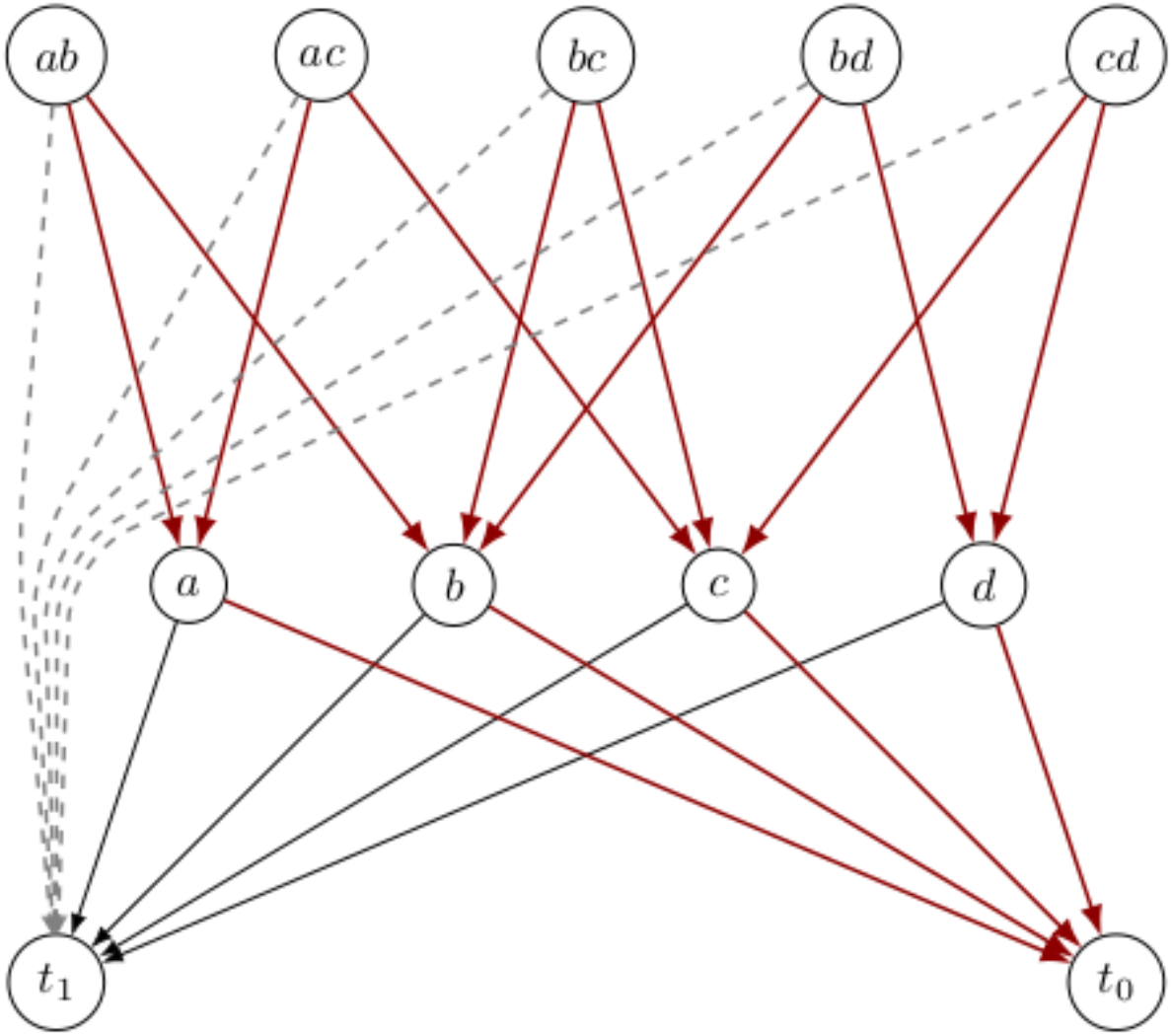
Constructed instance for *LBCA* with *eb* = |*E*| + k = 5 + 2 = 7. All thick red edges are forced edges since they are direct edges that are required, and the gray dashed edges are requirements where no edge exists. The excess branching of the red subgraph is already 5, and every internal vertex has an out-edge, which means we are allowed to add exactly two more edges. The only edges we can add are the edges that go from a vertex v ∈ {*a, b, c, d*) to t_1_, which will reveal a vertex cover in *G*.

## References

Aga, O. N., Brun, M., Giannakis, K., Dauda, K. A., Diaz-Uriarte, R., & Johnston, I. G. (2024). HyperTraPS-CT: Inference and prediction for accumulation pathways with flexible data and model structures. bioRxiv, 2024–03.

Allen, J. F. (2015). Why chloroplasts and mitochondria retain their own genomes and genetic systems: Colocation for redox regulation of gene expression. Proceedings of the National Academy of Sciences, 112(33), 10231–10238. 10.1073/pnas.1500012112

Angaroni, F., Chen, K., Damiani, C., Caravagna, G., Graudenzi, A., & Ramazzotti, D. (2022). PMCE: Efficient inference of expressive models of cancer evolution with high prognostic power. Bioinformatics, 38(3), 754–762.

Arbuckle, K., Bennett, C. M., & Speed, M. P. (2014). A simple measure of the strength of convergent evolution. Methods in Ecology and Evolution, 5(7), 685–693. 10.1111/2041-210X.12195

Bang-Jensen, J., & Gutin, G. (2018). Branching in Digraphs with Many and Few Leaves: Structural and Algorithmic Results. In B. Goldengorin (Ed.), Optimization Problems in Graph Theory: In Honor of Gregory Z. Gutin’s 60th Birthday (pp. 93– 106). Springer International Publishing. 10.1007/978-3-319-94830-0_5

Beerenwinkel, N., Däumer, M., Sing, T., Rahnenführer, J., Lengauer, T., Selbig, J., Hoffmann, D., & Kaiser, R. (2005). Estimating HIV evolutionary pathways and the genetic barrier to drug resistance. The Journal of Infectious Diseases, 191(11), 1953–1960.

Beerenwinkel, N., Eriksson, N., & Sturmfels, B. (2007). Conjunctive bayesian networks.

Beerenwinkel, N., Schwarz, R. F., Gerstung, M., & Markowetz, F. (2015). Cancer evolution: Mathematical models and computational inference. Systematic Biology, 64(1), e1–e25.

Björkholm, P., Harish, A., Hagström, E., Ernst, A. M., & Andersson, S. G. (2015). Mitochondrial genomes are retained by selective constraints on protein targeting. Proceedings of the National Academy of Sciences, 112(33), 10154– 10161. 10.1073/pnas.1421372112

Boyko, J. D., & Beaulieu, J. M. (2021). Generalized hidden Markov models for phylogenetic comparative datasets. Methods in Ecology and Evolution, 12(3), 468–478. 10.1111/2041-210X.13534

Boyko, J. D., O’Meara, B. C., & Beaulieu, J. M. (2023). A novel method for jointly modeling the evolution of discrete and continuous traits. Evolution, 77(3), 836–851. 10.1093/evolut/qpad002

Burger, G., Gray, M. W., Forget, L., & Lang, B. F. (2013). Strikingly Bacteria-Like and Gene-Rich Mitochondrial Genomes throughout Jakobid Protists. Genome Biology and Evolution, 5(2), 418–438. 10.1093/gbe/evt008

Butenko, A., Lukeš, J., Speijer, D., & Wideman, J. G. (2024). Mitochondrial genomes revisited: Why do different lineages retain different genes? BMC Biology, 22(1), 15. 10.1186/s12915-024-01824-1

Camin, J. H., & Sokal, R. R. (1965). A Method for Deducing Branching Sequences in Phylogeny. Evolution, 19(3), 311–326. 10.2307/2406441

Casali, N., Nikolayevskyy, V., Balabanova, Y., Harris, S. R., Ignatyeva, O., Kontsevaya, I., Corander, J., Bryant, J., Parkhill, J., Nejentsev, S., Horstmann, R. D., Brown, T., & Drobniewski, F. (2014). Evolution and transmission of drug-resistant tuberculosis in a Russian population. Nature Genetics, 46(3), Article 3. 10.1038/ng.2878

Catanzaro, D., Aringhieri, R., Di Summa, M., & Pesenti, R. (2015). A branch-price-and-cut algorithm for the minimum evolution problem. European Journal of Operational Research, 244(3), 753–765. 10.1016/j.ejor.2015.02.019

Chu, Y.-J. (1965). On the shortest arborescence of a directed graph. Scientia Sinica, 14, 1396–1400.

Dauda, K. A., Aga, O. N. L., & Johnston, I. G. (2024). Clustering large-scale biomedical data to model dynamic accumulation processes in disease progression and anti-microbial resistance evolution (p. 2024.09.19.613871). bioRxiv. 10.1101/2024.09.19.613871

Day, W. H. E., Johnson, D. S., & Sankoff, D. (1986). The computational complexity of inferring rooted phylogenies by parsimony. Mathematical Biosciences, 81(1), 33– 42. 10.1016/0025-5564(86)90161-6

Desper, R., Jiang, F., Kallioniemi, O.-P., Moch, H., Papadimitriou, C. H., & Schäffer, A. A. (1999). Inferring tree models for oncogenesis from comparative genome hybridization data. Journal of Computational Biology, 6(1), 37–51.

Diaz-Uriarte, R. (2023). A picture guide to cancer progression and monotonic accumulation models: Evolutionary assumptions, plausible interpretations, and alternative uses (2312.06824). arXiv. 10.48550/arXiv.2312.06824

Diaz-Uriarte, R., & Herrera-Nieto, P. (2022). EvAM-Tools: Tools for evolutionary accumulation and cancer progression models. Bioinformatics, 38(24), 5457– 5459.

Edmonds, J. (1967). Optimum branchings. Journal of Research of the National Bureau of Standards Section B Mathematics and Mathematical Physics, 71B(4), 233. 10.6028/jres.071B.032

El-Kebir, M. (2018). SPhyR: Tumor phylogeny estimation from single-cell sequencing data under loss and error. Bioinformatics, 34(17), i671–i679. 10.1093/bioinformatics/bty589

Farris, J. S. (1970). Methods for Computing Wagner Trees. Systematic Biology, 19(1), 83– 92. 10.1093/sysbio/19.1.83

Farris, J. S. (1977). Phylogenetic Analysis Under Dollo’s Law. Systematic Biology, 26(1), 77–88. 10.1093/sysbio/26.1.77

Felsenstein, J. (2003). Inferring Phylogenies. Oxford University Press.

Fernandes, L. M., & Gouveia, L. (1998). Minimal spanning trees with a constraint on the number of leaves. European Journal of Operational Research, 104(1), 250–261. 10.1016/S0377-2217(96)00327-X

Fernández-Baca, D. (2001). The Perfect Phylogeny Problem. In X.Z. Cheng & D.-Z. Du (Eds.), Steiner Trees in Industry (pp. 203–234). Springer US. 10.1007/978-1-4613-0255-1_6

Fernández-Baca, D., & Lagergren, J. (2003). A Polynomial-Time Algorithm for Near-Perfect Phylogeny. SIAM Journal on Computing, 32(5), 1115–1127. 10.1137/S0097539799350839

Fichera, M. E., & Roos, D. S. (1997). A plastid organelle as a drug target in apicomplexan parasites. Nature, 390(6658), 407–409. 10.1038/37132

Gao, Y., Gaither, J., Chifman, J., & Kubatko, L. (2022). A phylogenetic approach to inferring the order in which mutations arise during cancer progression. PLOS Computational Biology, 18(12), e1010560. 10.1371/journal.pcbi.1010560

Garey, M. R., & Johnson, D. S. (1979). Computers and intractability (Vol. 174). freeman San Francisco. https://bohr.wlu.ca/hfan/cp412/references/ChapterOne.pdf

Georgiadis, L. (2003). Arborescence optimization problems solvable by Edmonds’ algorithm. Theoretical Computer Science, 301(1), 427–437. 10.1016/S0304-3975(02)00888-5

Gerstung, M., Baudis, M., Moch, H., & Beerenwinkel, N. (2009). Quantifying cancer progression with conjunctive Bayesian networks. Bioinformatics, 25(21), 2809– 2815.

Giannakis, K., Arrowsmith, S. J., Richards, L., Gasparini, S., Chustecki, J. M., Røyrvik, E. C., & Johnston, I. G. (2022). Evolutionary inference across eukaryotes identifies universal features shaping organelle gene retention. Cell Systems, 13(11), 874-884.e5. 10.1016/j.cels.2022.08.007

Giannakis, K., Richards, L., & Johnston, I. G. (2024). Ecological predictors of organelle genome evolution: Phylogenetic correlations with taxonomically broad, sparse, unsystematized data. Systematic Biology, syae009. 10.1093/sysbio/syae009

Greenbury, S. F., Barahona, M., & Johnston, I. G. (2020). HyperTraPS: Inferring probabilistic patterns of trait acquisition in evolutionary and disease progression pathways. Cell Systems, 10(1), 39–51.

Gutin, G., Razgon, I., & Kim, E. J. (2008). Minimum Leaf Out-Branching Problems. In R. Fleischer & J. Xu (Eds.), Algorithmic Aspects in Information and Management (pp. 235–246). Springer. 10.1007/978-3-540-68880-8_23

Hjelm, M., Höglund, M., & Lagergren, J. (2006). New probabilistic network models and algorithms for oncogenesis. Journal of Computational Biology, 13(4), 853–865.

Hjort, K., Goldberg, A. V., Tsaousis, A. D., Hirt, R. P., & Embley, T. M. (2010). Diversity and reductive evolution of mitochondria among microbial eukaryotes. Philosophical Transactions of the Royal Society B: Biological Sciences, 365(1541), 713–727. 10.1098/rstb.2009.0224

Jahn, K., Kuipers, J., & Beerenwinkel, N. (2016). Tree inference for single-cell data. Genome Biology, 17(1), 86. 10.1186/s13059-016-0936-x

Johnston, I. G. (2019). Tension and Resolution: Dynamic, Evolving Populations of Organelle Genomes within Plant Cells. Molecular Plant, 12(6), 764–783. 10.1016/j.molp.2018.11.002

Johnston, I. G., & Diaz-Uriarte, R. (2024). A hypercubic Mk model framework for capturing reversibility in disease, cancer, and evolutionary accumulation modelling (p. 2024.06.27.600959). bioRxiv. 10.1101/2024.06.27.600959

Johnston, I. G., Hoffmann, T., Greenbury, S. F., Cominetti, O., Jallow, M., Kwiatkowski, D., Barahona, M., Jones, N. S., & Casals-Pascual, C. (2019). Precision identification of high-risk phenotypes and progression pathways in severe malaria without requiring longitudinal data. NPJ Digital Medicine, 2(1), 63.

Johnston, I. G., & Røyrvik, E. C. (2020). Data-driven inference reveals distinct and conserved dynamic pathways of tool use emergence across animal taxa. Iscience, 23(6).

Johnston, I. G., & Williams, B. P. (2016). Evolutionary Inference across Eukaryotes Identifies Specific Pressures Favoring Mitochondrial Gene Retention. Cell Systems, 2(2), 101–111. 10.1016/j.cels.2016.01.013

Karp, R. M. (2010). Reducibility Among Combinatorial Problems. In M. Jünger, T. M. Liebling, D. Naddef, G. L. Nemhauser, W. R. Pulleyblank, G. Reinelt, G. Rinaldi, & L. A. Wolsey (Eds.), 50 Years of Integer Programming 1958-2008 (pp. 219–241). Springer Berlin Heidelberg. 10.1007/978-3-540-68279-0_8

Keeling, P. J. (2010). The endosymbiotic origin, diversification and fate of plastids. Philosophical Transactions of the Royal Society B: Biological Sciences, 365(1541), 729–748. 10.1098/rstb.2009.0103

Kelly, S. (2021). The economics of organellar gene loss and endosymbiotic gene transfer. Genome Biology, 22(1), 345. 10.1186/s13059-021-02567-w

Krause, K. (2008). From chloroplasts to “cryptic” plastids: Evolution of plastid genomes in parasitic plants. Current Genetics, 54(3), 111–121. 10.1007/s00294-008-0208-8

Luo, X. G., Kuipers, J., & Beerenwinkel, N. (2023). Joint inference of exclusivity patterns and recurrent trajectories from tumor mutation trees. Nature Communications, 14(1), Article 1. 10.1038/s41467-023-39400-w

Lynch, M., Koskella, B., & Schaack, S. (2006). Mutation Pressure and the Evolution of Organelle Genomic Architecture. Science, 311(5768), 1727–1730. 10.1126/science.1118884

Mahapatra, S., Narayanan, M., & Narayanaswamy, N. S. (2024). Parameterized Algorithms for the Steiner Arborescence Problem on a Hypercube (2110.02830). arXiv. 10.48550/arXiv.2110.02830

Makiuchi, T., & Nozaki, T. (2014). Highly divergent mitochondrion-related organelles in anaerobic parasitic protozoa. Biochimie, 100, 3–17. 10.1016/j.biochi.2013.11.018

Moen, M. T., & Johnston, I. G. (2023). HyperHMM: Efficient inference of evolutionary and progressive dynamics on hypercubic transition graphs. Bioinformatics, 39(1), btac803.

Montazeri, H., Kuipers, J., Kouyos, R., Böni, J., Yerly, S., Klimkait, T., Aubert, V., Günthard, H. F., Beerenwinkel, N., & Study, S. H. C. (2016). Large-scale inference of conjunctive Bayesian networks. Bioinformatics, 32(17), i727–i735.

Morita, K., Wang, F., Jahn, K., Hu, T., Tanaka, T., Sasaki, Y., Kuipers, J., Loghavi, S., Wang, S. A., Yan, Y., Furudate, K., Matthews, J., Little, L., Gumbs, C., Zhang, J., Song, X., Thompson, E., Patel, K. P., Bueso-Ramos, C. E., … Takahashi, K. (2020). Clonal evolution of acute myeloid leukemia revealed by high-throughput single-cell genomics. Nature Communications, 11(1), 5327. 10.1038/s41467-020-19119-8

Nicol, P. B., Coombes, K. R., Deaver, C., Chkrebtii, O., Paul, S., Toland, A. E., & Asiaee, A. (2021). Oncogenetic network estimation with disjunctive Bayesian networks. Computational and Systems Oncology, 1(2), e1027.

Oborník, M., & Lukeš, J. (2015). The Organellar Genomes of Chromera and Vitrella, the Phototrophic Relatives of Apicomplexan Parasites. Annual Review of Microbiology, 69(Volume 69, 2015), 129–144. 10.1146/annurev-micro-091014-104449

O’Leary, N. A., Wright, M. W., Brister, J. R., Ciufo, S., Haddad, D., McVeigh, R., Rajput, B., Robbertse, B., Smith-White, B., Ako-Adjei, D., Astashyn, A., Badretdin, A., Bao, Y., Blinkova, O., Brover, V., Chetvernin, V., Choi, J., Cox, E., Ermolaeva, O., … Pruitt, K. D. (2016). Reference sequence (RefSeq) database at NCBI: Current status, taxonomic expansion, and functional annotation. Nucleic Acids Research, 44(D1), D733–D745. 10.1093/nar/gkv1189

Pagel, M. (1994). Detecting correlated evolution on phylogenies: A general method for the comparative analysis of discrete characters. Proceedings of the Royal Society of London. Series B: Biological Sciences, 255(1342), 37–45.

Peach, R. L., Greenbury, S. F., Johnston, I. G., Yaliraki, S. N., Lefevre, D. J., & Barahona, M. (2021). Understanding learner behaviour in online courses with Bayesian modelling and time series characterisation. Scientific Reports, 11(1), 2823.

Peterson, L. E., & Kovyrshina, T. (2017). Progression inference for somatic mutations in cancer. Heliyon, 3(4). 10.1016/j.heliyon.2017.e00277

Renz, J., Dauda, K. A., Aga, O. N. L., Diaz-Uriarte, R., Löhr, I. H., Blomberg, B., & Johnston, I. G. (2024). Evolutionary accumulation modelling in AMR: Machine learning to infer and predict evolutionary dynamics of multi-drug resistance (2411.00219). arXiv. 10.48550/arXiv.2411.00219

Roger, A. J., Muñoz-Gómez, S. A., & Kamikawa, R. (2017). The Origin and Diversification of Mitochondria. Current Biology: CB, 27(21), R1177–R1192. 10.1016/j.cub.2017.09.015

Ross, E. M., & Markowetz, F. (2016). OncoNEM: Inferring tumor evolution from single-cell sequencing data. Genome Biology, 17(1), 69. 10.1186/s13059-016-0929-9

Schill, R., Klever, M., Rupp, K., Hu, Y. L., Lösch, A., Georg, P., Pfahler, S., Vocht, S., Hansch, S., Wettig, T., Grasedyck, L., & Spang, R. (2024). Reconstructing Disease Histories in Huge Discrete State Spaces. KI - Künstliche Intelligenz. 10.1007/s13218-023-00822-9

Schill, R., Solbrig, S., Wettig, T., & Spang, R. (2020). Modelling cancer progression using mutual hazard networks. Bioinformatics, 36(1), 241–249.

Schwartz, R., & Schäffer, A. A. (2017). The evolution of tumour phylogenetics: Principles and practice. Nature Reviews Genetics, 18(4), 213–229. 10.1038/nrg.2016.170

Sloan, D. B., Alverson, A. J., Chuckalovcak, J. P., Wu, M., McCauley, D. E., Palmer, J. D., & Taylor, D. R. (2012). Rapid evolution of enormous, multichromosomal genomes in flowering plant mitochondria with exceptionally high mutation rates. PLoS Biology, 10(1), e1001241. 10.1371/journal.pbio.1001241

Sloan, D. B., Havird, J. C., & Sharbrough, J. (2017). The on-again, ob-again relationship between mitochondrial genomes and species boundaries. Molecular Ecology, 26(8), 2212–2236. 10.1111/mec.13959

Smith, D. R., & Keeling, P. J. (2015). Mitochondrial and plastid genome architecture: Reoccurring themes, but significant differences at the extremes. Proceedings of the National Academy of Sciences, 112(33), 10177–10184. 10.1073/pnas.1422049112

Speed, M. P., & Arbuckle, K. (2017). Quantification provides a conceptual basis for convergent evolution. Biological Reviews, 92(2), 815–829. 10.1111/brv.12257

Sridhar, S., Dhamdhere, K., Blelloch, G., Halperin, E., Ravi, R., & Schwartz, R. (2007). Algorithms for Efficient Near-Perfect Phylogenetic Tree Reconstruction in Theory and Practice. IEEE/ACM Transactions on Computational Biology and Bioinformatics, 4(4), 561–571. IEEE/ACM Transactions on Computational Biology and Bioinformatics. 10.1109/TCBB.2007.1070

Stayton, C. T. (2015). What does convergent evolution mean? The interpretation of convergence and its implications in the search for limits to evolution. Interface Focus, 5(6), 20150039. 10.1098/rsfs.2015.0039

Szabo, A., & Boucher, K. M. (2008). Oncogenetic trees. In Handbook of cancer models with applications (pp. 1–24). World Scientific.

von Heijne, G. (1986). Why mitochondria need a genome. FEBS Letters, 198(1), 1–4. 10.1016/0014-5793(86)81172-3

Williams, B. P., Johnston, I. G., Covshoff, S., & Hibberd, J. M. (2013). Phenotypic landscape inference reveals multiple evolutionary paths to C4 photosynthesis. eLife, 2, e00961. 10.7554/eLife.00961

Zhang, W., & Wang, S.-L. (2018). Inference of Cancer Progression With Probabilistic Graphical Model From Cross-Sectional Mutation Data. IEEE Access, 6, 22889–22898. IEEE Access. 10.1109/ACCESS.2018.2827024

